# Women’s brain aging: effects of sex-hormone exposure, pregnancies, and genetic risk for Alzheimer’s disease

**DOI:** 10.1101/826123

**Authors:** Ann-Marie G. de Lange, Claudia Barth, Tobias Kaufmann, Ivan I. Maximov, Dennis van der Meer, Ingrid Agartz, Lars T. Westlye

## Abstract

Sex hormones such as estrogen fluctuate across the female lifespan, with high levels during reproductive years and natural decline during the transition to menopause. Women’s exposure to estrogen may influence their heightened risk of Alzheimer’s disease (AD) relative to men, but little is known about how it affects normal brain aging. Recent findings from the UK Biobank demonstrate less apparent brain aging in women with a history of multiple childbirths, highlighting a potential link between sex-hormone exposure and brain aging. We investigated endogenous and exogenous sex-hormone exposure, genetic risk for AD, and neuroimaging-derived biomarkers for brain aging in 16,854 middle to older-aged women. The results showed that as opposed to parity, higher cumulative sex-hormone exposure was associated with more evident brain aging, indicating that i) high levels of cumulative exposure to sex-hormones may have adverse effects on the brain, and ii) beneficial effects of pregnancies on the female brain are not solely attributable to modulations in sex-hormone exposure. In addition, for women using hormonal replacement therapy (HRT), starting treatment earlier was associated with less evident brain aging, but only in women with a genetic risk for AD. Genetic factors may thus contribute to how timing of HRT initiation influences women’s brain aging trajectories.

## 1. Introduction

Women are at significantly greater risk of developing Alzheimer’s disease (AD) or other types of dementia relative to men (Laws et al., 2018). The genotype-related risk for developing AD is also modified by sex, with higher risk in female carriers of apolipoprotein E type 4 (APOE e4) compared to male carriers (Altmann et al., 2014). Emerging evidence suggests that APOE genotype may interact with effects of exogenous estrogen exposure (Yaffe, 2001; Srivastava et al., 1997; Stone et al., 1998), influencing dementia and AD risk (Yaffe et al., 2000). However, little is known about the influence of sex-hormone exposure on normal brain aging trajectories. Changes in sex hormones such as estradiol are known to influence brain plasticity (Galea et al., 2014; Simerly, 2002), and in premenopausal women, magnetic resonance imaging (MRI) studies have indicated modulating effects of endogenous estrogen fluctuations on brain structure across the menstrual cycle (Barth et al., 2016) and during pregnancy (Hoekzema et al., 2017). While higher endogenous estrogen levels have been associated with larger brain volumes during women’s reproductive years (Barth et al., 2016; Lisofsky et al., 2015), results from the Rotterdam Scan Study showed that in menopausal women, higher endogenous estrogen levels were associated with *smaller* brain volumes in specific areas (den Heijer et al., 2003). Negative effects of *exogenous* estrogen levels have also been reported, and findings from the Women’s Health Initiative Memory Study showed that conjugated estrogen, both alone and in combination with progestin, was associated with greater atrophy among women aged 65 years and older (Resnick et al., 2009). In addition, conjugated estrogen administration has been linked to higher rates of ventricular expansion over 4 years in recently menopausal women (Kantarci et al., 2016). However, other neuroimaging studies suggest a protective effect of hormone replacement therapy (HRT) on gray matter (Erickson et al., 2005), as well as white matter and ventricle size (Ha et al., 2007). Emerging evidence indicates that oral contraceptives (OC), another source of exogenous estrogen, affect aspects of brain structure and function in young adults (reviewed by (Pletzer & Kerschbaum, 2014)), but despite their widespread use (Christin-Maitre, 2013), the impact of OCs on brain aging is unknown.

We recently showed lower brain age in parous compared to nulliparous women in the UK Biobank cohort (de Lange et al., 2019). In the present paper, we investigate the association between estimates of sex-hormone exposure and brain aging beyond the effects of parity in 16,854 UK Biobank women (mean age 54.70 *±* 7.29 years). Brain-age prediction using machine learning and imaging-derived measures of cortical thickness, cortical volume, and subcortical volume (Fischl et al., 2002; Glasser et al., 2016; Kaufmann et al., 2019; de Lange et al., 2019) was performed to estimate ‘brain age’ (Franke & Gaser, 2019) for each participant. *Brain age gap*, calculated by subtracting chronological age from estimated brain age, was used as a measure of apparent brain aging (Franke & Gaser, 2019; Smith et al., 2019; Franke et al., 2010; Cole et al., 2017). Cumulative sex-hormone exposure was estimated by an *index of cumulative estrogen exposure (ICEE)* (Smith et al., 1999), including age at menarche and menopause, time since menopause, body mass index (BMI), and duration of HRT use. Exogenous exposure was estimated by HRT and OC use. To examine the effect of APOE e4 genotype on HRT usage and brain aging, we performed follow-up analyses including APOE e4 genotype interactions. As the ‘critical period hypothesis’ states that HRT may be neuroprotective if it is initiated close to menopause (MacLennan et al., 2006; Gibbs & Gabor, 2003), we further tested whether age at HRT initiation, both alone and in relation to age at menopause, was associated with apparent brain aging. In order to investigate the link between sex-hormone exposure and brain aging beyond the effects of parity (de Lange et al., 2019), all analyses were corrected for number of childbirths.

## 2. Materials and Methods

### 2.1. Sample

The sample was drawn from the UK Biobank (www.ukbiobank.ac.uk), and included 16,854 women. Sample demographics are provided in Tables 1, 2, and 3.

**Table 1:** Sample demographics. Ethnic background: W = white, B = black, M = mixed, A = Asian, C = Chinese, O = other. Educational qualification: U = university/college degree, A = A levels or equivalent, O = O levels/General Certificate of Secondary Education (GCSE) or equivalent, C = Certificate of Secondary Education (CSE) or equivalent, N = National Vocational Qualification (NVQ) or equivalent, P = professional qualification, e.g. nursing/teaching, Noa = none of the above. For the categories, see http://biobank.ctsu.ox.ac.uk/crystal/coding.cgi?id=100305 and http://biobank.ctsu.ox.ac.uk/crystal/coding.cgi?id=1001. Menopausal status was based on responses to the question *‘Have you had your menopause?’*. 10.50% answered ‘Not sure - had a hysterectomy’, 10.50% answered ‘Not sure - other reason’. N = sample size; SD = standard deviation. http://biobank.ndph.ox.ac.uk/showcase/field.cgi?id=2724.

|  |  |
| --- | --- |
| **N** | 16,854 |
| **Age [years] (mean ± SD)** | 54.70 ± 7.29 |
| **Ethnic background** | W 97.39 \| B 0.61 \| M 0.52 \| A 0.67 \| C 0.33 \| O 0.45 |
| **Education** | U 43.27 \| A 13.97 \| O 21.28 \| C 4.14 \| N 3.32 \| P 5.75 \| Noa 6.42 |
| **Menopausal status** | Yes 50.97% \| No 30.29% \| Not sure 21% \| Prefer not to answer 0.08% |

**Table 2:** Sample demographics for hormone replacement therapy (HRT) users (n = 5,651) and non-users (n = 11,172). Mean *±* standard deviation / % for each variable in each of the groups; N = sample size; BMI = body mass index.

|  | **HRT users** | **Non-users** | **t** | **$\chi^2$** | **p** | **N** |
| --- | --- | --- | --- | --- | --- | --- |
| **Age [years]** | 59.3 ± 5.4 | 52.4 ± 7.00 | 965.4 | | < 0.001 | 16,823 |
| **Number of births** | 1.8 ± 1.1 | 1.7 ± 1.2 | 144.1 | | < 0.001 | 16,817 |
| **Age at first birth [years]** | 25.7 ± 4.8 | 27.5 ± 5.1 | 600.8 | | < 0.001 | 13,270 |
| **BMI [kg/m<sup>2</sup>]** | 27.5 ± 4.8 | 27.4 ± 4.8 | 730.5 | | < 0.001 | 16,724 |
| **Age at menopause [years]** | 48.5 ± 5.8 | 50.6 ± 3.9 | 951.17 | | < 0.001 | 9,369 |
| **Menopause (yes)** | 96.1 % | 50.4 % |  | 2695.4 | < 0.001 | 14,179 |
| **Hysterectomy (yes)** | 31.5 % | 6.5 % |  | 1841.7 | < 0.001 | 16,797 |
| **Oophorectomy (yes)** | 17.6 % | 1.5 % |  | 1486.5 | < 0.001 | 16,712 |
| **Hypertension (yes)** | 21.6 % | 16.2% |  | 51.0 | < 0.001 | 12,029 |

**Table 3:**
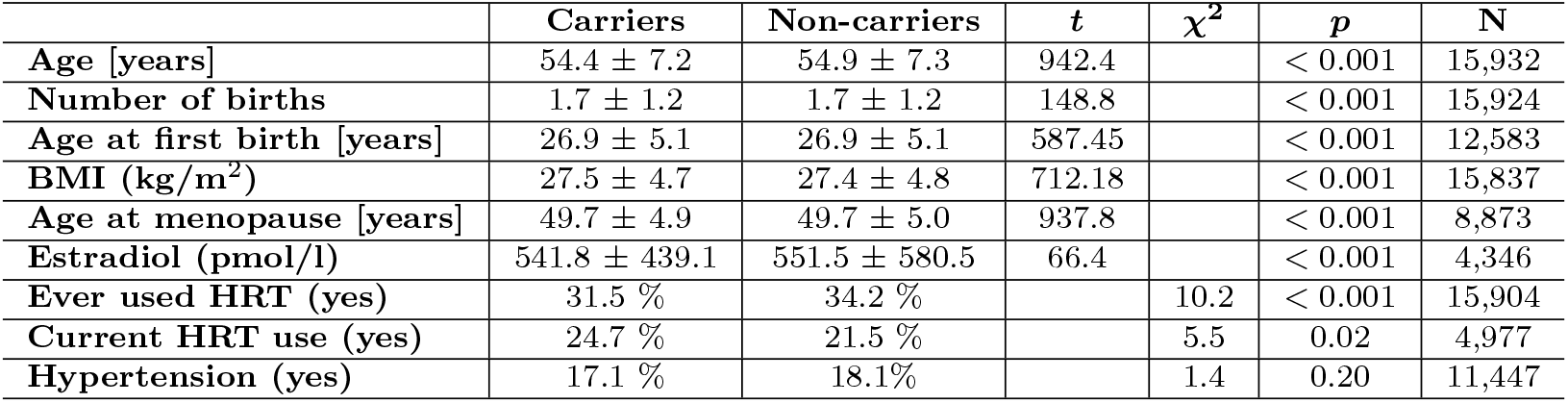
Sample demographics for apolipoprotein E type 4 (APOE e4) genotype carriers (n = 4,277) and non-carriers (n = 11,655). Mean *±* standard deviation / % for each variable in each of the groups; N = sample size; BMI = body mass index; HRT = hormone replacement therapy.

|  | **Carriers** | **Non-carriers** | **t** | **$\chi^2$** | **p** | **N** |
| --- | --- | --- | --- | --- | --- | --- |
| **Age [years]** | 54.4 ± 7.2 | 54.9 ± 7.3 | 942.4 | | < 0.001 | 15,932 |
| **Number of births** | 1.7 ± 1.2 | 1.7 ± 1.2 | 148.8 | | < 0.001 | 15,924 |
| **Age at first birth [years]** | 26.9 ± 5.1 | 26.9 ± 5.1 | 587.45 | | < 0.001 | 12,583 |
| **BMI (kg/m<sup>2</sup>)** | 27.5 ± 4.7 | 27.4 ± 4.8 | 712.18 | | < 0.001 | 15,837 |
| **Age at menopause [years]** | 49.7 ± 4.9 | 49.7 ± 5.0 | 937.8 | | < 0.001 | 8,873 |
| **Estradiol (pmol/l)** | 541.8 ± 439.1 | 551.5 ± 580.5 | 66.4 | | < 0.001 | 4,346 |
| **Ever used HRT (yes)** | 31.5 % | 34.2 % |  | 10.2 | < 0.001 | 15,904 |
| **Current HRT use (yes)** | 24.7 % | 21.5 % |  | 5.5 | 0.02 | 4,977 |
| **Hypertension (yes)** | 17.1 % | 18.1% |  | 1.4 | 0.20 | 11,447 |

### 2.2. Sex-hormone exposure

Women who had missing data, or had responded ‘do not know’ or ‘prefer not to answer’ for any of the relevant variables, were excluded for each analysis. For relevant analyses, women with a reported age at menarche (n = 1) or age at OC start (n = 1) at 5 years were excluded. ICEE was approximated by including age at menarche and menopause, duration of HRT in years, BMI, and time since menopause in years. The variables were first standardized and then either added (duration HRT, age at menopause, BMI) to or subtracted (age at menarche, time since menopause) from the index depending on their impact on endogenous estrogen (Smith et al., 1999). After removing women with missing data, 8,878 women were included in the ICEE analyses involving the brain-age model. For HRT, 11,139 never-users and 5,546 users were included in the analyses. For OC, 2,213 never-users and 14,615 users were included in the analyses. Women who had never used HRT/OC were coded 0, while current and former users were coded 1. Multiple regression analyses were run to investigate the association between each estimate of sex-hormone exposure and brain age gap. All analyses were corrected for number of childbirths and age. In addition, hysterectomy and/or oophorectomy were included as covariates in the HRT models, and current HRT use, ever used HRT, and length since menopause in years were included a covariates in the APOE e4 status × circulating estradiol level models. To test whether other known confounders including education, BMI, hypertensive status, and age at first birth could influence the results, additional models including these variables were run. Hypertensive status (yes/no) was defined as systolic blood pressure ≥ 140 mmHg and diastolic blood pressure ≥ 90 mmHg, otherwise individuals were classified as non-hypertensive (Warren et al., 2017). The statistical analyses were conducted using R, version 3.5.2, and Python 3.

### 2.3. Genotyping

For genotyping, we used the UK Biobank version 3 imputed data, which has undergone extensive quality control procedures as described by the UK Biobank genetics team (Bycroft et al., 2018). The APOE e genotype was approximated based on the two APOE e single-nucleotide polymorphisms - rs7412 and rs429358 (Lyall et al., 2016). Further information on the genotyping process is available in the UK Biobank documentation (www.ukbiobank.ac.uk/scientists-3/genetic-data), including detailed technical documentation (genotyping_workflow.pdf). APOE e4 status was labelled *carrier* for e3/e4 and e4/e4 combinations, and *non-carrier* for e2/e2, e2/e3 and e3/e3 combinations (Lyall et al., 2019). The homozygous e2/e4 allele combination was removed due to its ambiguity with e1/e3 (Wisdom et al., 2011).

### 2.4. Hormone assay

Serum blood samples were taken at day of MRI scan. Estradiol was analyzed at the UK Biobank’s purpose-build laboratory in Stockport, and measured by two step competitive analysis on a Unicel DXI 800 Access Immunoassay System (Beckman Coulter, UK, Ltd; analytical range: 73 - 17621 pmol/l). Further information on the immunoassay and quality control steps is available in the UK Biobank documentation (serum_biochemistry.pdf). To investigate whether APOE e4 status interacted with circulating estradiol levels in menopausal women, a multiple linear regression was run including an *APOE e4 status × circulating estradiol level* interaction term. The model was corrected for current HRT use, ever used HRT, length since menopause, and number of births.

### 2.5. MRI processing

A detailed overview of the data acquisition, protocol parameters, and image validation can be found in (Alfaro-Almagro et al., 2018) and (Miller et al., 2016). Raw T1-weighted MRI data for all participants were processed using a harmonized analysis pipeline, including automated surface-based morphometry and subcortical segmentation as implemented in FreeSurfer 5.3 (Fischl et al., 2002). In line with recent large-scale implementations (Kauf-mann et al., 2019; de Lange et al., 2019), we utilized a fine-grained cortical parcellation scheme (Glasser et al., 2016) to extract cortical thickness, area, and volume for 180 regions of interest per hemisphere, in addition to the classic set of subcortical and cortical summary statistics from FreeSurfer (Fischl et al., 2002). This yielded a total set of 1,118 structural brain imaging features (360/360/360/38 for cortical thickness/area/volume, as well as cerebellar/subcortical and cortical summary statistics, respectively). The MRI variables were residualized with respect to scanning site, ethnic background, intracranial volume, and Freesurfer-derived Euler numbers (Rosen et al., 2018) using linear models. To remove outliers, participants with Euler numbers of standard deviation (SD) *±* 4 were identified and excluded (n = 159). In addition, participants with SD *±* 4 on the global MRI measures mean cortical or subcortical gray matter volume were excluded (n = 79 and n = 13, respectively), yielding a total of 16,854 participants with T1-weighted MRI data.

### 2.6. Principal component analysis (PCA)

A PCA was run with z-transformed MRI variables *z* = (*x* — *μ*)/σ, where *x* is an MRI variable of mean *μ* and standard deviation σ). The top 100 components were used in the subsequent analyses, explaining 56.48% of the total variance. As a cross check, the relationship between ICEE and brain age gap was re-analyzed with 200 components, explaining 70.61% of the total variance. With 200 components included, the association between ICEE and brain age gap was *β =* 0.03, *SE =* 0.01, *t =* 2.42*,p =* 0.02. As the results were consistent, 100 components were chosen to reduce computational time.

### 2.7. Brain age prediction

The XGBRegressor model from XGBoost (https://xgboost.readthedocs.io/en/latest/python/index.html) was used to run the brain age prediction analysis with an algorithm that has been used in recent large-scale brain age studies (de Lange et al., 2019; Kaufmann et al., 2019; Smith et al., 2019). Parameters were set to max depth = 3, number of estimators = 100, and learning rate = 0.1 (defaults). The predicted age based on the PCA components was estimated in a 10-fold cross validation, assigning an estimated brain age value to each individual. Brain age gap was calculated using (estimated brain age - chronological age). Average RMSE and R^2^ were calculated from a 10-fold cross validation with 10 repetitions per fold, and compared to null distributions calculated from 10,000 permutations. The results are shown in Figure 1.

**Figure 1:**
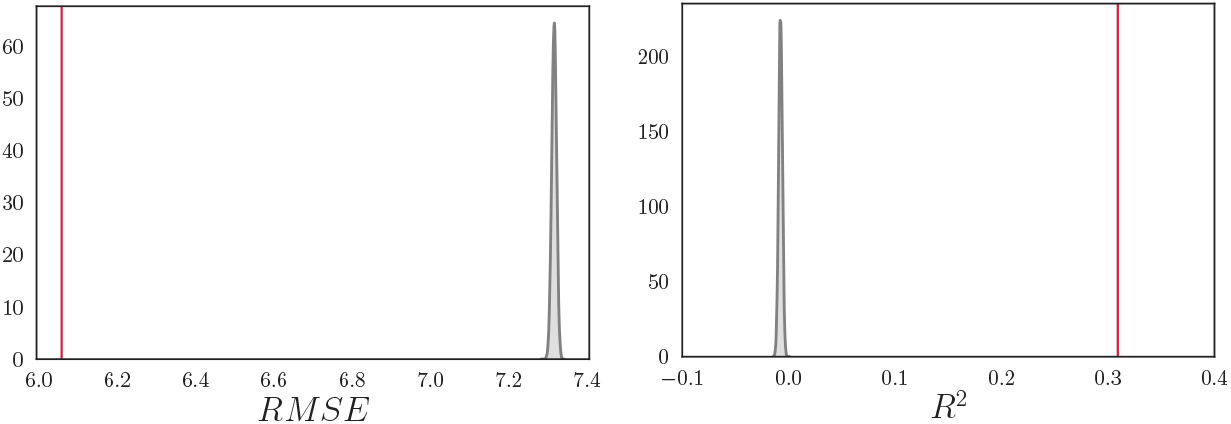
Average root mean square error (RMSE) and R^2^ compared to null distributions. Left: The mean *±* standard deviation (SD) RMSE was 6.06 *±* 0.09, based on a 10-fold cross validation with 10 repetitions per fold (red vertical line). The null distribution calculated from 10,000 permutations is shown in gray, with a mean *±* SD of 7.32 *±* 0.006. The number of permuted results from the null distribution that exceeded the mean from the cross validation was 0 (p = 1.00 × 10^-4^). Right: The mean *±* SD R^2^ for the brain age model was 0.31 *±* 0.09 (red vertical line). The null distribution calculated from 10,000 permutations is shown in gray, with a mean *±* SD of —0.007 *±* 0.002 (p =1.00 × 10^-4^).

## 3. Results

The accuracy of the brain-age prediction model is shown in Table 4. The associations between estimates of sex-hormone exposure and apparent brain aging are summarized in Table 5 and Figure 2; p-values are reported before and after false discover rate correction (*p_corr_*) (Benjamini & Hochberg, 1995).

**Table 4:**
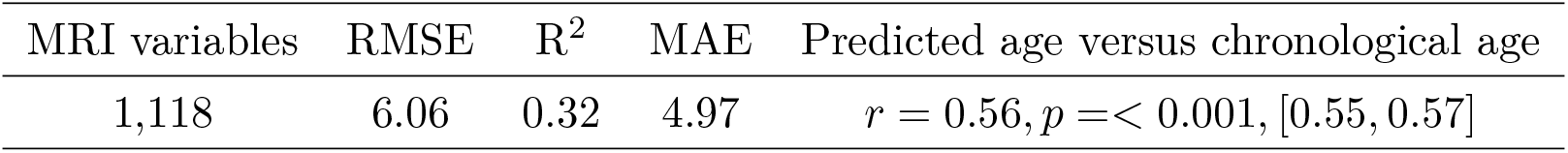
Number of magnetic resonance imaging (MRI) variables, root mean square error (RMSE), R^2^, mean absolute error (MAE), and the correlation between predicted and chronological age. RMSE and MAE are reported in years. 95% confidence intervals are indicated in square brackets.

| MRI variables | RMSE | $R^2$ | MAE | Predicted age versus chronological age |
| --- | --- | --- | --- | --- |
| 1,118 | 6.06 | 0.32 | 4.97 | $r = 0.56, p = < 0.001, [0.55, 0.57]$ |

**Table 5:** The associations between each estimate of sex-hormone exposure and apparent brain aging. β = slope, SE = standard error. ICEE = Index of cumulative estrogen exposure. Hormone replacement therapy (HRT) and oral contraceptive (OC) status = 0 for never-users and 1 for current and former users. Each analysis included age and number of births as a covariates, in addition to specific covariates for each measure as detailed in Section 3. P-values are reported before and after false discovery rate (FDR)-correction (*P_corr_*) (Benjamini & Hochberg, 1995). Corrected p-values below 0.05 are marked with an asterisk.

| ICEE |  |  |  |  |
| --- | --- | --- | --- | --- |
| $\beta$ | SE | $t$ | $p$ | $p_{corr}$ |
| 0.040 | 0.014 | 2.809 | 0.005 | 0.018* |
| HRT status |  |  |  |  |
| $\beta$ | SE | $t$ | $p$ | $p_{corr}$ |
| 0.169 | 0.055 | 3.073 | 0.002 | 0.015* |
| Age at HRT initiation |  |  |  |  |
| $\beta$ | SE | $t$ | $p$ | $p_{corr}$ |
| 0.022 | 0.009 | 2.509 | 0.012 | 0.028* |
| Duration of HRT usage |  |  |  |  |
| $\beta$ | SE | $t$ | $p$ | $p_{corr}$ |
| 0.004 | 0.005 | 0.787 | 0.432 | 0.503 |
| OC status |  |  |  |  |
| $\beta$ | SE | $t$ | $p$ | $p_{corr}$ |
| 0.017 | 0.066 | 0.250 | 0.802 | 0.802 |

### 3.1. Index of cumulative estrogen exposure (ICEE)

A multiple linear regression showed a positive association between ICEE and apparent brain aging as shown in Table 5 and Figure 2, indicating that when correcting for number of previous childbirths and age, higher ICEE was linked to more apparent brain aging (n = 8,878). The inclusion of education, hypertensive status, and age at first birth as covariates yielded similar results (*β* = 0.045, *SE* = 0.019, *t* = 2.380, *p* = 0.017, *p_corr_* = 0.030, n = 4,791). The associations between apparent brain aging and reproductive span, calculated as (age at menopause — age at menarche), as well as age at menarche and menopause separately, are provided in the Supplementary Information (SI). Differential effects of surgical vs. natural menopause on brain aging were also examined (see SI).

**Figure 2:**
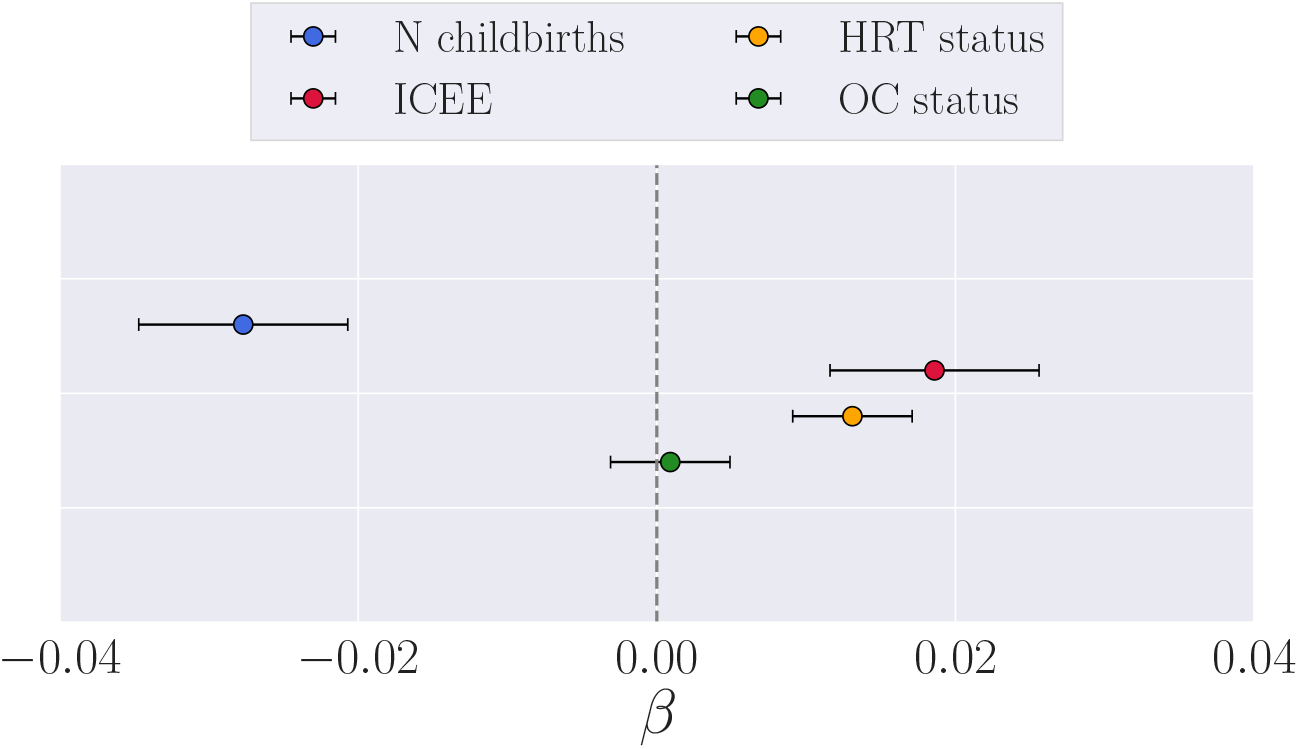
Associations between estimates of sex-hormone exposure and apparent brain aging. ICEE = index of cumulative estrogen exposure. The points show the *β* values (slope) from separate multiple regression analyses with brain age gap (see Section 2.7 in Materials and Methods) as dependent variable, and *number of births* (covariates: age and ICEE), *ICEE* (covariates: age and number of births), *hormone replacement therapy (HRT) status* (covariates: age, number of births, had hysterectomy and/or oophorectomy), and *oral contraceptive (OC) status* (covariates: age, number of births) as independent variables. To obtain a direct comparison of *β* values in the plot, all variables were standardized prior to performing the multiple regressions (subtracting the mean and dividing by the SD). HRT and OC status = 0 for never-users and 1 for current and former users. The error bars represent the standard error on the *β*.

### 3.2. Exogenous sex-hormone exposure

A positive association was found between *hormone replacement therapy (HRT) status* and apparent brain aging in pre-menopausal and menopausal women, with less evident brain aging in never-users (n = 11,139) compared to users (n = 5,546 with 1,182 still using; covariates: number of births, had hysterectomy and/or oophorectomy, and age). When including age at first birth, education, BMI, and hypertensive status in the model, the results were similar (β = 0.168, *SE* = 0.074, *t* = 2.262, *p* = 0.024, *p_corr_* = 0.033, never-users = 6,236, n users = 2,971). No significant associations were found between *Oral contraceptive (OC) status* and apparent brain aging, as shown in Figure 2 and Table 5 (covariates: number of births and age; number of never-users vs users was 2,213 vs 14,615. 398 women were still using OC).

Within the group of HRT users (n = 5,164), a positive relationship was found between *age at HRT initiation* and apparent brain aging as shown in Table 5, indicating less evident brain aging in women who started HRT treatment earlier (covariates: number of births, had hysterectomy and/or oophorectomy, and age). No significant association was found between *duration of HRT use* (age last used HRT - age started HRT) and apparent brain aging (covariates: number of births, age, had hysterectomy and/or oophorectomy, n = 5,164).

### 3.3. APOE eĄ genotype and HRT initiation

In HRT users, there was an effect of *APOE eĄ status × age started HRT* on apparent brain aging, as shown in Table 6, with a trend-level association after applying FDR correction (*p_corr_* = 0.063). Follow-up analyses showed that (1) the relationship between age at HRT initiation and apparent brain aging was confined to the carrier group (n = 1,227), as shown in Figure 3, and that (2) in menopausal APOE e4 carriers, *age for HRT onset relative to age at menopause* (age at menopause – age started HRT) was positively associated with apparent brain aging, indicating beneficial effects of HRT initiation before onset of menopause (β = —0.078, SE = 0.023, *t* = — 3.441, *p* = 6.083 × *10^-4^,p_corr_* = 0.003, n = 826, covariates: number of births, age, had hysterectomy and/or oophorectomy). There was no dose-dependent effect of the interaction age started HRT × carrier group (n with 1 e4 allele = 1,126, n with 2 e4 alleles =101, *β* = 0.085, *SE* = 0.064, *t* = 1.317, *p* = 0.188, *p_corr_* = 0.188).

**Figure 3:**
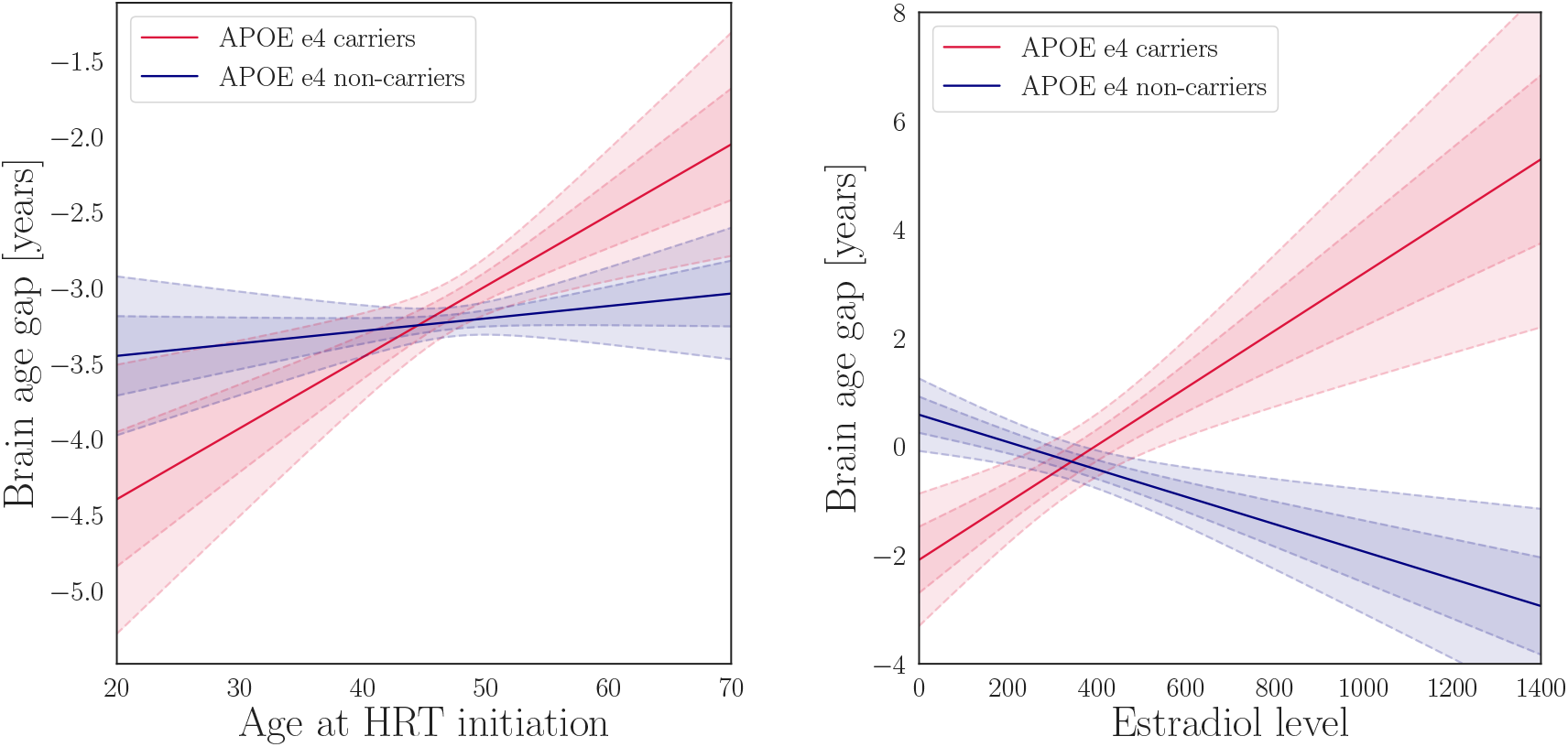
Apolipoprotein E (APOE) genotype interactions. Left plot: The lines show the association (β) between age started hormone replacement therapy (HRT) and apparent brain aging for the APOE e4 carriers (red) and non-carriers (blue). The fitted values are corrected for the covariates in the model (age, number of births, had hysterectomy, and/or oophorectomy). Right plot: The lines show the association between estradiol levels and apparent brain aging for the APOE e4 carriers (red) and non-carriers (blue). The fitted values are corrected for the covariates in the model (age, number of births, current HRT use, ever used HRT, length since menopause, age at first birth, and education). The shaded areas show the 68.3% (1 SD) and 95% (2 SD) confidence intervals for each fit.

**Table 6:**
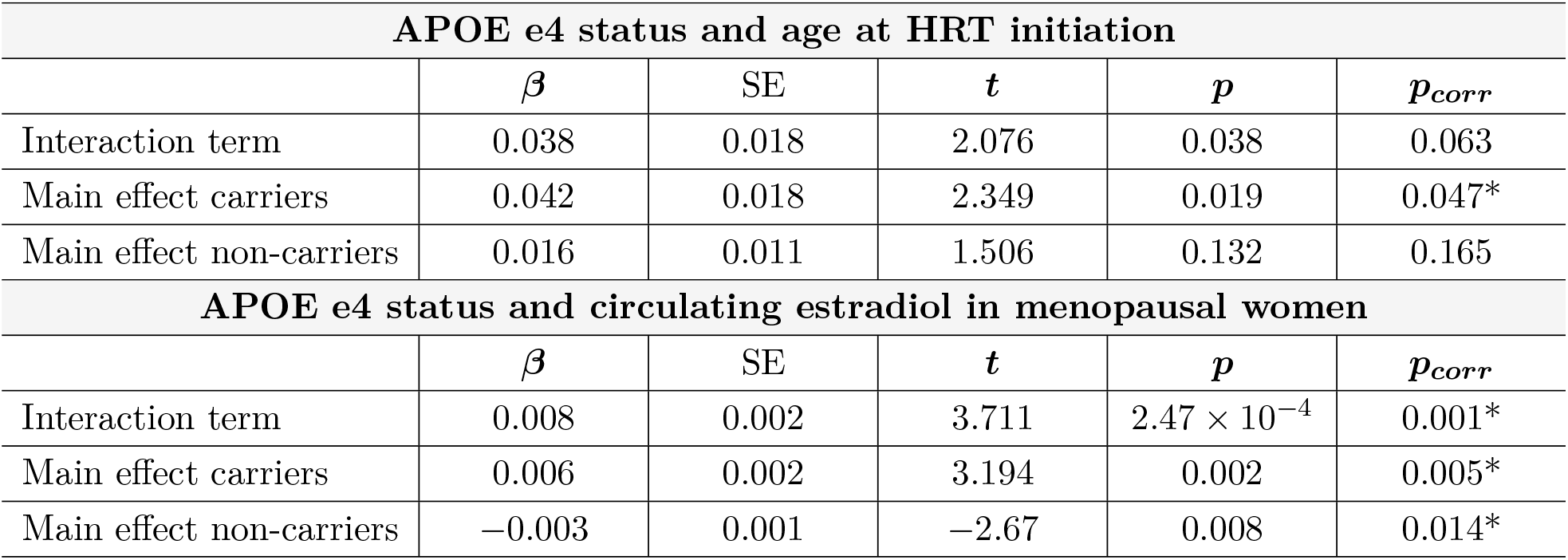
The associations between apolipoprotein E type 4 (APOE e4) × age at hormone replacement therapy (HRT) initiation and APOE e4 × circulating estradiol, and main effects within carrier and non-carriers. *β* = slope, SE = standard error. P-values are reported before and after false discovery rate correction (*p_corr_*) (Benjamini & Hochberg, 1995). Corrected p-values below 0.05 are marked with an asterisk.

| APOE e4 status and age at HRT initiation |  |  |  |  |  |
| --- | --- | --- | --- | --- | --- |
| | $\beta$ | SE | $t$ | $p$ | $p_{corr}$ |
| Interaction term | 0.038 | 0.018 | 2.076 | 0.038 | 0.063 |
| Main effect carriers | 0.042 | 0.018 | 2.349 | 0.019 | 0.047* |
| Main effect non-carriers | 0.016 | 0.011 | 1.506 | 0.132 | 0.165 |
| APOE e4 status and circulating estradiol in menopausal women |  |  |  |  |  |
| | $\beta$ | SE | $t$ | $p$ | $p_{corr}$ |
| Interaction term | 0.008 | 0.002 | 3.711 | $2.47 \times 10^{-4}$ | 0.001* |
| Main effect carriers | 0.006 | 0.002 | 3.194 | 0.002 | 0.005* |
| Main effect non-carriers | -0.003 | 0.001 | -2.67 | 0.008 | 0.014* |

### 3.4. APOE e4 genotype and circulating estradiol levels

As shown in Figure 3, significant cross-over interaction effects of *APOE e4 status × circulating estradiol levels* on apparent brain aging were found in menopausal women (β = 0.003, *SE* = 0.001, *t* = 2.270, *p* = 0.024, *p_corr_* = 0.030, n = 539, covariates: current HRT use, ever used HRT, length since menopause, and number of births). When including age at first birth, education, hypertensive status, and BMI as covariates, the results showed a stronger effect of the interaction, as well as a negative main effect of estradiol levels on apparent brain aging (β = —0.003, *SE* = 0.001, *t* = — 2.699, *p* = 0.007). Based on this finding, the subsequent follow-up analyses were also corrected for education, age at birth, hypertensive status, and BMI. A positive main effect of estradiol levels on apparent brain aging was found in APOE e4 carriers (n = 101), while a negative main effect was found in non-carriers (n = 310), as shown in Figure 3. No main association was found between APOE e4 status and apparent brain aging (β = 0.049, *SE* = 0.051, *t* = 0.954, *p* = 0.340, *p_corr_* = 0.340, covariates: number of births and age; number of carriers vs non-carriers = 4,276 vs 11,649).

The analyses were re-run with corrections for (1) neurological and mental conditions, and (2) polygenic risk for Alzheimer’s disease. The results are provided in the SI. In brief, these corrections did not influence the results.

## 4. Discussion

The results showed that as opposed to parity, higher cumulative and exogenous sex-hormone exposure was associated with more evident brain aging. *In vitro* studies have shown that exposure to low concentrations of 17β-estradiol promoted neuronal survival, whereas exposure to high concentrations was ineffective and led to increased cellular susceptibility to neurodegenerative insults (Chen et al., 2006). It has also been demonstrated that low-dose estrogen replacements have anti-inflammatory properties, whereas higher dosages show increases in inflammatory markers such as C-reactive protein (Prestwood et al., 2004). In line with this, the current findings indicate that high levels of exposure to sex-hormones may have adverse effects on women’s brain aging.

While estrogen levels rise up to 300-fold during pregnancy (Schock et al., 2016), they fall 100-1000 fold postnatally (Nott et al., 1976), and parous women have shorter menstrual cycles and lower levels of estradiol than nulliparous women (Bernstein et al., 1985; Dorgan et al., 1995). Hormonal modulations contribute to maternal brain adaptations in pregnancy and postpartum (Galea et al., 2014; Kinsley & Lambert, 2008), but their long-term effects on brain aging are not fully understood. The present results indicate that beneficial effects of pregnancies on brain aging may relate to factors beyond sex-hormone fluctuations. Other pregnancy-related mechanisms including immune regulations (Mor et al., 2011; Hillerer et al., 2014; Luppi, 2003) may have implications for inflammatory susceptibility later in life (Fox et al., 2018; Natri et al., 2019), subsequently impacting women’s brain aging trajectories. For instance, one study found that higher cumulative time spent pregnant in first trimesters, but not third trimesters, conferred a protective effect against AD (Fox et al., 2018), indicating that immune processes such as the the proliferation of regulatory T cells (Kieffer et al., 2017), which is highest in the first trimester, could be more relevant for AD risk relative to estrogen exposure. In line with this, the ‘pregnancy compensation hypothesis’ suggests that pregnancies involves long-lasting, favorable regulations of the female immune system (Natri et al., 2019), which could underlie observed differences in apparent brain aging between parous and nulliparous women (de Lange et al., 2019).

The cessation of ovarian hormone function during menopause has been linked to altered inflammatory processes involving increases in cytokine levels, and changes in T cell biology (reviewed by (Mishra & Brinton, 2018)). For some, these processes may constitute a menopausal immune senescence that may increase the risk for AD (Fox et al., 2018; Wyss-Coray & Rogers, 2012). The ‘critical period hypothesis’ states that HRT may be neuroprotective if it is initiated near the time of cessation of ovarian function - approximately within 5 years of menopause (MacLennan et al., 2006; Gibbs & Gabor, 2003). Our results lend further support to this hypothesis, as earlier age at HRT initiation, particularly before menopause, was associated with less evident brain aging. However, this relationship was present in APOE e4 carriers only, indicating that genetic factors may contribute to how timing of HRT initiation influences women’s brain aging trajectories. Higher menopausal levels of estradiol were linked to more evident brain aging in carriers, while lower estradiol levels was linked to more evident brain aging in non-carriers. In line with this, increased estradiol levels induced by estrogen therapy have been associated with reduced risk of developing AD in APOE e4 non-carriers, but not in carriers (Yaffe et al., 2000; Manly et al., 2000).

To the best of our knowledge, the current work is the first comprehensive study of the associations between endogenous and exogenous hormone exposure, APOE genotype, and normal brain aging in a population-based cohort. Large-scale population-based studies enable the identification of subtle effects that could go undetected in smaller samples, and are key to foster understanding of factors contributing to brain-aging processes and risk for neurodegenerative disease. However, the cross-sectional nature of the presented data does not enable causal inference, and longitudinal studies are required to fully understand how sex-hormone exposure influences women’s brain health across the lifespan. Furthermore, the cohort data lacks details on HRT and OC formulation, administration, and dosage, and different compound compositions and modes of administration may affect brain aging differently. The lack of information on breastfeeding, which is known to reduce cumulative exposure to endogenous estrogen (Bernstein, 2002), may also influence the precision of our ICEE approximation.

An issue with genetic aging studies is the bias towards survivors (Heffernan et al., 2016), which implies that the number of APOE e4 carriers could be lower in older-age cohorts. In the present study, age was negatively associated with number of carriers (*r* = —0.03, *p* =< 0.001, 95%CI = [—0.05, —0.02]), indicating potential survival-bias in the sample. Further, estradiol levels were only available for a subset of women, and within this subset, a relatively high proportion (80%) had oestradiol values in the lower range. The UK Biobank notes that this reflects the menopausal status of the participants at recruitment (with 25% being premenopausal; see https://biobank.ctsu.ox.ac.uk/crystal/crystal/docs/biomarker_issues.pdf), which is expected given the age range of the cohort. Based on these aspects, it should be noted that the presented results may not apply to populations beyond those represented in UK Biobank (Haworth et al., 2019).

In conclusion, our study provides evidence of an association between higher sex-hormone exposure and more apparent brain aging, indicating that i) high levels of exposure to sex-hormones may have adverse effects on the brain, and ii) beneficial effects of pregnancies relate to factors beyond sex-hormone fluctuations. Further, the influence of sex-hormone trajectories on the brain may be genotype-specific, with more prominent effects of timing and dosage of hormone replacements in women with a genetic risk for AD. These findings represent an important contribution to the understudied field of female-specific factors and women’s brain aging (Galea et al., 2018), which may complement prospective longitudinal studies on women’s brain health and epidemiological sex-differences in AD.

## Acknowledgments

This research was conducted using the UK Biobank under Application 27412, and received funding from the Research Council of Norway (286838, 273345, 249795, 276082), and the European Research Council (ERC) under the European Union’s Horizon 2020 research and innovation programme (Grant agreement No. 802998). The permutation testing was performed using resources provided by UNINETT Sigma2 - the National Infrastructure for High Performance Computing and Data Storage in Norway. All authors declare that they have no conflict of interest.

## Author contributions

A-M.G.dL., C.B., T.K., and L.T.W. designed the study; A-M.G.dL., C.B., T.K., I.M. and D.vdM. performed the data analysis; A-M.G.dL., and C.B. interpreted the data as well as drafted and finalized the manuscript. A-M.G.dL. and C.B. contributed equally to the work. I.A., L.T.W., T.K, I.I.M. and D.vdM critically revised the first draft and approved the final manuscript.

## 5. Supplementary Information

*P*-values are reported before and after false discovery rate (FDR)-correction.

### 1. Correction for ICD-10 diagnoses

To evaluate whether disorders known to affect the brain could drive the observed effects, all the main analyses were rerun after excluding participants with the following main and secondary ICD-10 diagnoses: F (Mental and behavioral disorder, n = 84), G (Diseases of the nervous system, n = 211) and I60-I69 (Cerebrovascular diseases, n = 42). Five women had the same or overlapping diagnoses for both main and secondary ICD-10 diagnosis.

In summary, the main results were not influenced by excluding participants with ICD-10 diagnoses. The corrected associations with apparent brain aging were as follows: *ICEE*: *β* = 0.035, *SE* = 0.014, *t* = 2.481, *p* = 0.013, *p_corr_* = 0.022, n = 8,701; *HRT status: β* = 0.169, *SE* = 0.055, *t* = 3.043, *p* = 0.002, *p_corr_* = 0.012, n = 16,358, user = 5450, never-user = 10908; *age started HRT*: *β* = 0.022, *SE* = 0.009,*t* = 2.476, *p* = 0.013, *p_corr_* = 0.022, n = 5,077; *duration of HRT use: β* = 0.004, *SE* = 0.005, *t* = 0.755, *p* = 0.450, *p_corr_* = 0.563, n = 5,077; and *OC status: β* = 0.031, *SE* = 0.067, *t* = 0.464, *p* = 0.643, *p_corr_* = 0.643, n users = 14,333, never-users = 2,168. The interaction of *APOE e4 status × estradiol levels* also yielded equivalent results:β = 0.008, *SE* = 0.002, *t* = 3.580, *p* = 4.062 × 10^-4^, *p_corr_* = 0.001, n = 289, covariates: current HRT use, ever used HRT, length since menopause, number of births, age at first birth, education, BMI and hypertensive status). The same was true for the main effect of estradiol levels on brain age in APOE e4 carriers *β* = 0.006, *SE* = 0.002, *t* = 3.090, *p* = 0.003, *p_corr_* = 0.004, n = 75, and non-carriers *β* = —0.003, *SE* = 0.001,*t* = —2.519, *p* = 0.013, *p_corr_* = 0.013, n = 214.

### 2. Correction for polygenic risk score (PRS) of Alzheimer’s disease

To examine whether polygenic risk for Alzheimer’s disease could drive the observed effects, all the main analyses were rerun while accounting for individual PRS scores. Individual PRS were calculated using PRSice version 1.25 (Euesden et al., 2014) at a p-value threshold of 0.05, using PRSice default settings. This includes the removal of the major histocompatibility complex (MHC; chromosome 6, 26-33Mb) and thinning of SNPs based on linkage disequilibrium (LD) and p-value. We based the PRS on Lambert and colleagues work (Lambert et al., 2013). No associations were found between PRS and apparent brain aging (β = 330.873, SE = 192.759, *t* = 330.873, *p* = 0.086, *p_corr_* = 0.129).

The main results were not influenced by PGRS scores. The corrected associations with apparent brain aging were as follows: *ICEE*: *β* = 0.040, *SE* = 0.014, *t* = 2.795, *p* = 0.005, *p_corr_* = 0.016, n = 8,618; *HRT status: β* = 0.167,SE = 0.056, *t* = 3.003, *p* = 0.003, *p_corr_* = 0.016, n = 16,177, user = 5,368, never-user = 10,809; *age started HRT*: *β* = 0.022, *SE* = 0.009,*t* = 2.423, *p* = 0.015, *p_corr_* = 0.031, n = 5,000; *duration of HRT use: β* = 0.004, *SE* = 0.005, *t* = 0.812, *p* = 0.418, *p_corr_* = 0.500, n = 5,000; and *OC status:* β = −0.007,*SE* = 0.067,*t* = −0.098, *p* = 0.922, *p_corr_* = 0.922, n = 16,314, user = 14,165, never-user = 2,148.

### 3. Reproductive span

Reproductive span was calculated as (age at menopause — age at menarche). 9,188 menopausal women had data on both variables and were included in the analysis. A multiple linear regression including number of births, ever used HRT, ever used OC, and age as covariates showed a trend towards a positive association between reproductive span and apparent brain aging, but the effect did not survive FDR correction (β = 0.012, *SE* = 1.872, *t* = 1.872, *p* = 0.061, *p_corr_* = 0.209).

### 4. Age at menarche

A multiple linear regression including number of births and age showed a negative association between age at menarche and apparent brain aging in pre- and menopausal women (β = —0.039, *SE* = 0.014, *t* = —2.751, *p* = 5.955 × 10^-3^, *p_corr_* = 0.054, n = 16,435), indicating more apparent brain aging with earlier age at menarche. However, when including age at first birth, education, BMI and hypertensive status as additional covariates, the association did not persist (*β* = —0.035, *SE* = 0.019, *t* = —1.814, *p* = 0.070, *p_corr_* = 0.209, n = 9,107).

### 5. Age at menopause

When accounting for a history of hysterectomy and/or oophorectomy, number of births and age, we found no relationship between age at menopause and apparent brain aging in menopausal women (β = 0.007,*SE* = 0.007, *t* = 0.976, *p* = 0.329, *p_corr_* = 0.429, n = 9,346).

### 6. Surgical vs. natural menopause

Surgical menopause is defined by women transitioning to menopause through removal of both ovaries (bilateral oophorectomy) rather than natural reproductive aging. Functioning ovaries can also be removed at time of hysterectomy to reduce the risk of ovarian cancer. We stratified menopausal women according to (1) natural menopause (n = 7,888) defined by absence of hysterectomy and/or oophorectomy, and (2) surgical menopause (n = 422) characterized by age at hysterectomy and/or oophorectomy coinciding with age at menopause. A linear regression including number of births, age, ever used HRT and time since menopause showed no association between type of menopause and apparent brain aging (*β* = —0.150, *SE* = 0.155, *t* = —0.967, *p* = 0.334, *p_corr_* = 0.429).

