## Supplementary Information for "Women’s brain aging: effects of sex-hormone exposure, pregnancies, and genetic risk for Alzheimer’s disease" (from another version of this preprint: version 1, 2019-10-31)

To examine whether APOE e4 status influenced the observed associations between estrogen exposure and brain aging, we performed linear models including an interaction term for APOE e4 status for each of the main analyses showing significant results, including the same covariates as provided in section 2.1 - 2.3. The main results are summarized in SI Table 1.

**SI Table 1:** The associations between APOE e4 status and brain age gap, and APOE e4 status * each of the estimates of hormone exposure and brain age gap based on the full T1w model. c = carriers, nc = non-carriers. P-values are reported before and after FDR correction. $SE$ = standard error.

| **Main effects: APOE e4 status and brain age gap**, c = 4276, nc = 11649 | | | | | |
|---|---|---|---|---|---|
| Model | $\beta$ | $SE$ | $t$ | $p$ | $p_{corr}$ |
| Full T1w model | 0.05 | 0.05 | 1.07 | 0.28 | 0.55 |
| Cortical thickness | 0.03 | 0.05 | 0.55 | 0.58 | 0.80 |
| Cortical volume | $-0.01$ | 0.04 | $-0.25$ | 0.80 | 0.88 |
| Subcortical volume | 0.10 | 0.06 | 1.73 | 0.08 | 0.33 |
| Diffusion | 0.04 | 0.07 | 0.52 | 0.61 | 0.80 |
| **Index of cumulative estrogen exposure * APOE e4 status (ICEE)**, c = 2208, nc = 6198 | | | | | |
| Model | $\beta$ | $SE$ | $t$ | $p$ | $p_{corr}$ |
| Full T1w model | 0.02 | 0.03 | 0.51 | 0.61 | 0.80 |
| Cortical thickness | 0.04 | 0.03 | 1.26 | 0.21 | 0.55 |
| Cortical volume | $-0.03$ | 0.03 | $-1.17$ | 0.24 | 0.55 |
| Subcortical volume | 0.04 | 0.04 | 1.06 | 0.29 | 0.55 |
| Diffusion | 0.05 | 0.05 | 1.19 | 0.23 | 0.55 |
| **Reproductive span * APOE e4 status**, c = 2287, nc = 6408 | | | | | |
| Model | $\beta$ | $SE$ | $t$ | $p$ | $p_{corr}$ |
| Full T1w model | 0.01 | 0.01 | 0.67 | 0.50 | 0.80 |
| Cortical thickness | 0.03 | 0.01 | 2.00 | 0.05 | 0.21 |
| Cortical volume | $5.47 \times 10^{-4}$ | 0.01 | 0.05 | 0.96 | 0.96 |
| Subcortical volume | 0.01 | 0.02 | 0.39 | 0.07 | 0.84 |
| Diffusion | 0.01 | 0.02 | 0.30 | 0.76 | 0.86 |
| **Hormone replacement therapy (HRT) status * APOE e4 status**, c = 4238, nc = 11536 | | | | | |
| Model | $\beta$ | $SE$ | $t$ | $p$ | $p_{corr}$ |
| Full T1w model | 0.01 | 0.11 | 0.07 | 0.95 | 0.96 |
| Cortical thickness | 0.04 | 0.11 | 0.33 | 0.74 | 0.86 |
| Cortical volume | $-0.01$ | 0.10 | $-0.1$ | 0.92 | 0.96 |
| Subcortical volume | 0.16 | 0.12 | 1.26 | 0.21 | 0.55 |
| Diffusion | 0.01 | 0.15 | 0.66 | 0.51 | 0.80 |
| **Age at HRT onset * APOE e4 status**, c = 1227, nc = 3646 | | | | | |
| Model | $\beta$ | $SE$ | $t$ | $p$ | $p_{corr}$ |
| Full T1w model | 0.04 | 0.02 | 2.18 | 0.03 | 0.18 |
| Cortical thickness | 0.05 | 0.02 | 2.66 | 0.01 | 0.07 |
| Cortical volume | 0.02 | 0.02 | 1.12 | 0.26 | 0.55 |
| Subcortical volume | 0.03 | 0.02 | 1.37 | 0.17 | 0.55 |
| Diffusion | 0.03 | 0.02 | 1.11 | 0.27 | 0.55 |
| **age started HRT relative to menopause * APOE e4 status**, c = 1227, nc = 3646 | | | | | |
| Model | $\beta$ | $SE$ | $t$ | $p$ | $p_{corr}$ |
| Full T1w model | $-0.08$ | 0.02 | $-3.34$ | $8.80 \times 10^{-4}$ | 0.02 |
| Cortical thickness | $-0.06$ | 0.02 | $-2.38$ | 0.02 | 0.13 |
| Cortical volume | $-0.04$ | 0.02 | $-2.00$ | 0.05 | 0.21 |
| Subcortical volume | $-0.03$ | 0.03 | $-1.3$ | 0.2 | 0.55 |
| Diffusion | $-0.03$ | 0.03 | $-0.84$ | 0.4 | 0.72 |

**Correction for ICD-10 diagnoses:**
To evaluate whether disorders known to affect the brain could drive the observed effects, all the main analyses were rerun after excluding the following main and secondary ICD-10 diagnoses: F (Mental and behavioral disorder, n = 84), G (Diseases of the nervous system, n = 211) and I60-I69 (Cerebrovascular diseases, n = 42). Five women had the same or overlapping diagnoses for both main and secondary ICD-10 diagnosis. In summary, the main results were not influenced by excluding participants with ICD-10 diagnoses. The corrected results for brain age gap based on the full T1w model were as follows: **ICEE**: $\beta = 0.04, SE = 0.01, t = 3.09, p = 2.02 \times 10^{-3}$, n = 8,701; **reproductive span:** $\beta = 0.02, SE = 5.92 \times 10^{-3}, t = 3.00, p = 2.74 \times 10^{-3}$, n = 9,005; **HRT status**: $\beta = 0.16, SE = 0.05, t = 3.25, p = 1.15 \times 10^{-3}$, n = 16,358, user = 5450, never-user = 10908; **age started HRT**: $\beta = 0.02, SE = 8.16 \times 10^{-3}, t = 2.74, p = 6.22 \times 10^{-3}$, n = 5,077; **duration of HRT use**: $\beta = 3.27 \times 10^{-3}, SE = 4.85 \times 10^{-3}, t = 0.68, p = 0.50$, n = 5,077; and **OC status**: $\beta = 0.11, SE = 0.07, t = 1.62, p = 0.11$, n users = 14,333, never-users = 2,168; ***in OC users***: **age started OC**: $\beta = 4.00 \times 10^{-3}, SE = 5.78 \times 10^{-3}, t = 0.69, p = 0.49$, n = 14,004; and **duration of OC use**: $\beta = 6.91 \times 10^{-3}, SE = 3.11 \times 10^{-3}, t = 2.22, p = 0.03$, n = 13,207.An ICD-10 corrected linear regression with a interaction of APOE e4 status * estradiol levels yielded in similar results: $\beta = 7.42 \times 10^{-3}, SE = 1.75 \times 10^{-3}, t = 4.24, p = 2.79 \times 10^{-5}$, n = 399, covariates: current HRT use, ever used HRT, length since menopause, number of births, age at first birth, education). The same was true for the main effect of estradiol levels on brain age based on the full T1w model in APOE e4 carriers $\beta = 5.11 \times 10^{-3}, SE = 1.55 \times 10^{-3}, t = 3.30, p = 1.41 \times 10^{-3}$, n = 98, and non-carriers $\beta = -2.60 \times 10^{-3}, SE = 8.57 \times 10^{-4}, t = -3.04, p = 2.63 \times 10^{-3}$, n = 298.

**Correction for polygenic risk score (PRS) of Alzheimer's disease:**
To examine whether polygenic risk for Alzheimer's disease could drive the observed effects, all the main analyses were rerun while accounting for individual PRS scores. Individual PRS were calculated using PRSice version 1.25 (**?** ) at a p-value threshold of 0.05, using PRSice default settings. This includes the removal of the major histocompatibility complex (MHC; chromosome 6, 26-33Mb) and thinning of SNPs based on linkage disequilibrium (LD) and p-value. We based the PRS on Lambert and colleagues work (**?** ). No associations were found between PRS and brain age gap (full T1w model:$\beta = 342.91, SE = 192.29, t = 1.78, p = 0.08$; cortical thickness: $\beta = 265.08, SE = 200.59, t = 1.32, p = 0.19$; cortical Volume: $\beta = 119.43, SE = 169.52, t = 0.71, p = 0.48$; subcortical volume: $\beta = 290.89, SE = 218.51, t = 1.33, p = 0.18$; diffusion: $\beta = 242.72, SE = 257.98, t = 0.94, p = 0.35$).

The main results were not influenced by PGRS scores. The corrected results for brain age gap based on the full T1w model were as follows: **ICEE**: $\beta = 0.05, SE = 0.01, t = 3.42, p = 6.35 \times 10^{-4}$, n = 8,618; **reproductive span:** $\beta = 0.02, SE = 6.00 \times 10^{-3}, t = 3.18, p = 1.49 \times 10^{-3}$, n = 8,917; **HRT status**: $\beta = 0.16, SE = 0.05, t = 3.21, p = 1.35 \times 10^{-3}$, n = 16,177, user = 5,368, never-user = 10,809; **age started HRT**: $\beta = 0.02, SE = 8.21 \times 10^{-3}, t = 2.59, p = 9.52 \times 10^{-3}$, n = 5,000; **duration of HRT use**: $\beta = 3.35 \times 10^{-3}, SE = 4.87 \times 10^{-3}, t = 0.69, p = 0.49$, n = 5,000; and **OC status**: $\beta = 0.07, SE = 0.07, t = 1.05, p = 0.29$, n = 16,314, user = 14,165, never-user = 2,148; **age started OC**: $\beta = 4.81 \times 10^{-4}, SE = 5.82 \times 10^{-3}, t = 0.08, p = 0.93$, n = 13,846; and **duration of OC use**: $\beta = 7.07 \times 10^{-3}, SE = 3.12 \times 10^{-3}, t = 2.26, p = 0.02$, n = 13,055.

**Surgical vs. natural menopause:**
Surgical menopause is defined by women transitioning to menopause through removal of both ovaries (bilateral oophorectomy) rather than natural reproductive aging. Functioning ovaries can also be removed at time of hysterectomy to reduce the risk of ovarian cancer. We stratified menopausal women according to (1) natural menopause (n = 7,888) defined by absence of hysterectomy and/or oophorectomy, and (2) surgical menopause (n = 422) characterized by age at hysterectomy and/or oophorectomy coinciding with age at menopause. A linear regression showed no association between type of menopause and brain age gap based on the full T1w model ($\beta = -0.17, SE = 0.15, t = -1.11, p = 0.27$, covariates: number of births, ever used HRT, time since menopause), or any of the other sub-models (cortical thickness: $\beta = -0.26, SE = 0.16, t = -1.65, p = 0.10$; cortical volume: $\beta = -0.05, SE = 0.13, t = -0.41, p = 0.68$; subcortical volume: $\beta = -0.17, SE = 0.17, t = -0.98, p = 0.32$; diffusion: $\beta = -0.22, SE = 0.19, t = -1.11, p = 0.27$, natural n = 4,535, surgical n = 282). Additional adjustment for age at first birth and education, did not influence the results (full T1w model:$\beta = -0.18, SE = 0.17, t = -1.10, p = 0.27$, natural n = 6,272, surgical n = 356). Due to the negative results, no follow-up analyses were conducted.

**Age at menarche:**
A multiple linear regression including number of births showed a negative association between age at menarche and brain age gap based on the full T1w model in pre- and menopausal women ($\beta = -0.04, SE = 0.01, t = -2.98, p = 2.93 \times 10^{-3}$, n = 16,435), indicating higher brain age relative to chronological age with earlier age at menarche. When including age at first birth and education as additional covariates, the results were similar ($\beta = -0.04, SE = 0.02, t = -2.33, p = 0.02$, n = 12,769). Significant relationships were found between age at menarche and brain age based on cortical thickness: $\beta = -0.04, SE = 0.02, t = -2.73, p = 6.37 \times 10^{-3}$), cortical volume: $\beta = -0.05, SE = 0.01, t = -4.06, p = 4.89 \times 10^{-5}$,

and subcortical volume $\beta = -0.05, SE = 0.02, t = -3.16, p = 1.60 \times 10^{-3}$).No significant association was found between age at menarche and brain age gap based on diffusion ($\beta = -0.01, SE = 0.02, t = -0.69, p = 0.49$, n = 9,561).

**Age at menopause:**

While accounting for a history of hysterectomy and/or oophorectomy as well as number of births, we found a positive relationship between age at menopause and brain age gap based on the full T1w model in menopausal women ($\beta = 0.02, SE = 6.48 \times 10^{-3}, t = 2.28, p = 0.02$, n = 9,346), indicating that an older age at menopause could be associated with a higher brain age gap. This association became trend-level significant after adjusting for age at first birth and education ($\beta = 0.01, SE = 7.39 \times 10^{-3}, t = 1.8, p = 0.07$, n = 7,431). No significant relationships were found between age at menopause and brain age gap based on cortical thickness ($\beta = 0.01, SE = 6.67 \times 10^{-3}, t = 1.83, p = 0.07$) or diffusion (n = 5,382,$\beta = -2.08 \times 10^{-3}, SE = 0.01, t = -0.24, p = 0.81,$), but significant, positive relationships were found between age at menopause and brain age gap based on subcortical ($\beta = 0.02, SE = 7.33 \times 10^{-3}, t = 2.60, p = 9.28 \times 10^{-3}$) and cortical volume ($\beta = 0.01, SE = 5.68 \times 10^{-3}, t = 2.24, p = 0.03$).
